# Comment on “Rapid and efficient *in vivo* astrocyte-to-neuron conversion with regional identity and connectivity?”

**DOI:** 10.1101/2020.09.02.279273

**Authors:** Gong Chen, Wen Li, Zongqin Xiang, Liang Xu, Minhui Liu, Qingsong Wang, Wenliang Lei

## Abstract

Regenerating functional new neurons in the adult mammalian central nervous system (CNS) has been proven to be very challenging due to the inability of neurons to divide and repopulate themselves after neuronal loss. In contrast, glial cells in the CNS can divide and repopulate themselves under injury or disease conditions. Therefore, many groups around the world have been able to utilize internal glial cells to directly convert them into neurons for neural repair. We have previously demonstrated that ectopic expression of NeuroD1 in dividing glial cells can directly convert reactive glial cells into neurons. However, Wang et al. recently posted an article in bioRxiv challenging the entire field of *in vivo* glia-to-neuron conversion after using one single highly toxic dose of AAV (2×10^13^ gc/ml, 1 μl) in the mouse cortex, producing artifacts that are very difficult to interpret. We present data here that reducing AAV dosage to safe level will avoid artifacts caused by toxic dosage. We also demonstrate with Aldh1l1-CreER^T2^ and Ai14 reporter mice that lineage-traced astrocytes can be successfully converted into NeuN^+^ neurons after infected by AAV5 GFAP::NeuroD1. Retroviral expression of NeuroD1 further confirms our previous findings that dividing glial cells can be converted into neurons. Together, the incidence of Wang et al. sends an alarming signal to the entire *in vivo* reprogramming field that the dosage of viral vectors is a critical factor to consider when designing proper experiments. For AAV, we recommend a relatively safe dose of 1×10^10^ - 1×10^12^ gc/ml (~1 μl) in the rodent brain for cell conversion experiments addressing basic science questions. For therapeutic purpose under injury or diseased conditions, AAV dosage needs to be adjusted through a series of dose finding experiments. Moreover, we recommend that the AAV results are further verified with retroviruses that mainly express transgenes in dividing glial cells in order to draw solid conclusions.

## INTRODUCTION

The idea of directly converting one type of cells into other cell types for tissue regeneration has fascinated biologists for decades (1). However, it was not until the pioneering research on master regulator genes that started to provide important insights on lineage reprogramming. For instance, transcription factor MyoD can convert dermal fibroblasts, chondroblasts and retinal pigmented epithelial cells into contracting muscle cells (2–4). Similarly, transcription factor C/EBP reprograms B lymphocytes into macrophages (5), while transcription factor Math1 transforms non-sensory cells into hair cells in the ear (6, 7). Besides that, neural transcription factor NeuroD converts most of the embryonic ectoderm cells into neurons in *Xenopus* (8). The cell transdifferentiation field entered into a new era led by the success of Shinya Yamanaka and his colleagues showing successful reprogramming of fibroblast cells into induced pluripotent stem cells (iPSCs) (9–11). In particular, using combinations of transcription factors and small molecules, many labs around the globe have been able to directly convert different types of cells into neurons both *in vitro* and *in vivo*. For example, Vierbuchen et al. converted skin fibroblast cells into neurons using transcription factors Ascl1, Brn2, and Myt1l (12). Shortly after that, many somatic cells such as fibroblasts, hepatocytes, pericytes, astrocytes, and peripheral T cells in cell culture have also been successfully trans-differentiated into various subtypes of induced neurons including but not limited to glutamatergic, GABAergic, dopaminergic, motor neurons, and retinal neurons *in vitro* (13–30). As for *in vivo* reprogramming, our group has previously reported that a single neural transcription factor NeuroD1 can convert reactive glial cells into fully functional neurons in mouse brains with injury or Alzheimer’s disease (31). More recently, we demonstrated that NeuroD1 AAV-based gene therapy can regenerate and protect a large number of functional neurons to restore brain functions after ischemic injury in adult mice (32). We also reported that AAV-mediated expression of NeuroD1 and Dlx2 can reprogram striatal astrocytes into GABAergic medium spiny neurons and hence improve the motor functions and extend the life span in Huntington’s disease mouse models (33). In another attempt to reprogram glial cells into neurons, Zhang and colleagues converted astrocytes into neuroblasts with transcription factor Sox2 and then further differentiated them into neurons in mouse brain and spinal cord (34–38). Many other groups have also successfully transdifferentiated glial cells into neurons *in vivo* through ectopic expression of Ascl1 (39) or combinations of transcription factors such as Ascl1+Lmx1a+Nurr1 (40, 41), or Ascl1+Sox2 (42), or Neurogenin-2+Bcl-2 (43), or Neurogenin-2 plus growth factors FGF2 and EGF (44). A mixture of NeuroD1, Ascl1, Lmx1a, and microRNA 218 transformed mouse astrocytes into dopaminergic neurons (45). In addition, overexpression of Ascl1 in mouse retina also converted Müller glia into inner retinal neurons in both young and adult mice with NMDA damage (46, 47), and application of Otx2, Crx and Nrl after β-catenin expression could reprogram Müller glia into rod photoreceptors which restored lost vision in adult mice (48). Different from overexpression of transcription factors, Qian et al. reported recently that depleting the RNA-binding protein Ptbp1 in the substantia nigra can convert midbrain astrocytes into dopaminergic neurons and restore motor functions in Parkinson’s disease mouse model (49). Surprisingly, Zhou et al. reported that striatal astrocytes can also be converted into dopaminergic neurons by CRISPR-mediated Ptbp1 knockdown (50), which has been disputed by Qian et al. (49). Taken together, direct glia-to-neuron conversion has been successfully achieved both *in vitro* and *in vivo*, using a variety of neural transcription factors or knocking down RNA-binding protein Ptbp1 by many labs around the world. Therefore, it is a completely surprise that Wang et al. (51) would challenge the entire field of *in vivo* glia-to-neuron conversion simply based on a set of experiments using very high dose of AAV at 2×10^13^ gc/ml (1 μl) in the mouse cortex. This article will attempt to clarify the confusion about the leakage versus conversion caused by highly toxic level of AAV used by Wang et al. (51). We also present evidence that lineage traced-astrocytes can be converted into neurons by NeuroD1 in Aldh1l1-CreER^T2^ mice crossed with Ai14 mice, and that using retrovirus to express transgenes in dividing glial cells is another safeguard to unambiguously demonstrate *in vivo* glia-to-neuron conversion.

## RESULTS

### AAV GFAP::Cre should express Cre in astrocytes not in neurons

It is perhaps not too difficult to understand that injecting too much viruses into the brain will cause toxic effects. However, it appears that not everyone knows the importance of viral dosing, or even worse, not checking whether there are any toxic effects after viral injection into the brain. A perfect example is the recent bioRxiv paper posted by Wang et al (51), where 2×10^13^ gc/ml × 1 μl AAV particles were injected into the mouse cortex, producing artifacts that led the authors to challenge the entire field of *in vivo* reprogramming.

AAV has been approved by FDA for various clinical trials due to its relatively low immunogenicity, and some gene therapy products based on AAV have been marketed for therapeutic use. In the gene therapy field, it is well-known that AAV dosing is critical when considering how much AAV should be administered into the body. For the brain, it is even more important to use minimal effective dosing of AAV to avoid any brain damage. Previously, it has been reported that high dosing AAV can produce harmful effects on both neurons and glial cells in mammalian brains (52–57). In particular, Ortinski et al (52) has reported that high titre AAV will cause astrocytic gliosis and impair synaptic transmission. Xiong et al (58) has also reported AAV toxicity in the retina when using much lower dose than that used by Wang et al (51). Unfortunately, Wang et al (51) appeared to be unaware of these very important works in the field and conducted all their experiments based on a single toxic dosing of AAV (2×10^13^ gc/ml × 1 μl) in the mouse cortex. Wang et al reported that when they injected AAV GFAP::Cre into the mouse brain, which should express Cre in GFAP^+^ astrocytes, they instead observed Cre expression predominantly in neurons (Wang et al., Fig. 3, 14 days post viral injection) (51). Typically, when one sees such abnormal result, one would immediately lower the AAV dosing and repeat the experiments until find the right dosing so that GFAP::Cre is properly expressed in GFAP^+^ astrocytes. However, it is surprising that the authors continued their experiments with such high level of AAV which is toxic to the CNS as reported before (52–57). With such high dosing of AAV, it is not surprising that Cre, and likely other transgenes as well such as NeuroD1, would be found in neurons, making any data interpretation invalid.

Such artifacts caused by high dosing of AAV reported by Wang et al (51) can be easily avoided using lower dosage of AAV. In fact, we have performed many GFAP::Cre experiments and never observed such high expression of Cre transgene in neurons, because we usually use much lower dosing AAV to express Cre (1×10^10^ - 1×10^11^ gc/ml × 1 μl). Fig. 1 illustrates a typical example of Cre expression in astrocytes (top row, GFAP/S100b^+^), but not in neurons (bottom row, NeuN^+^). Therefore, it is critical to design experiments properly with the right dosage of AAV to start any experiments. Too high dosage of AAV will damage brain cells and produce artifacts that is difficult to interpret.

**Figure 1.**
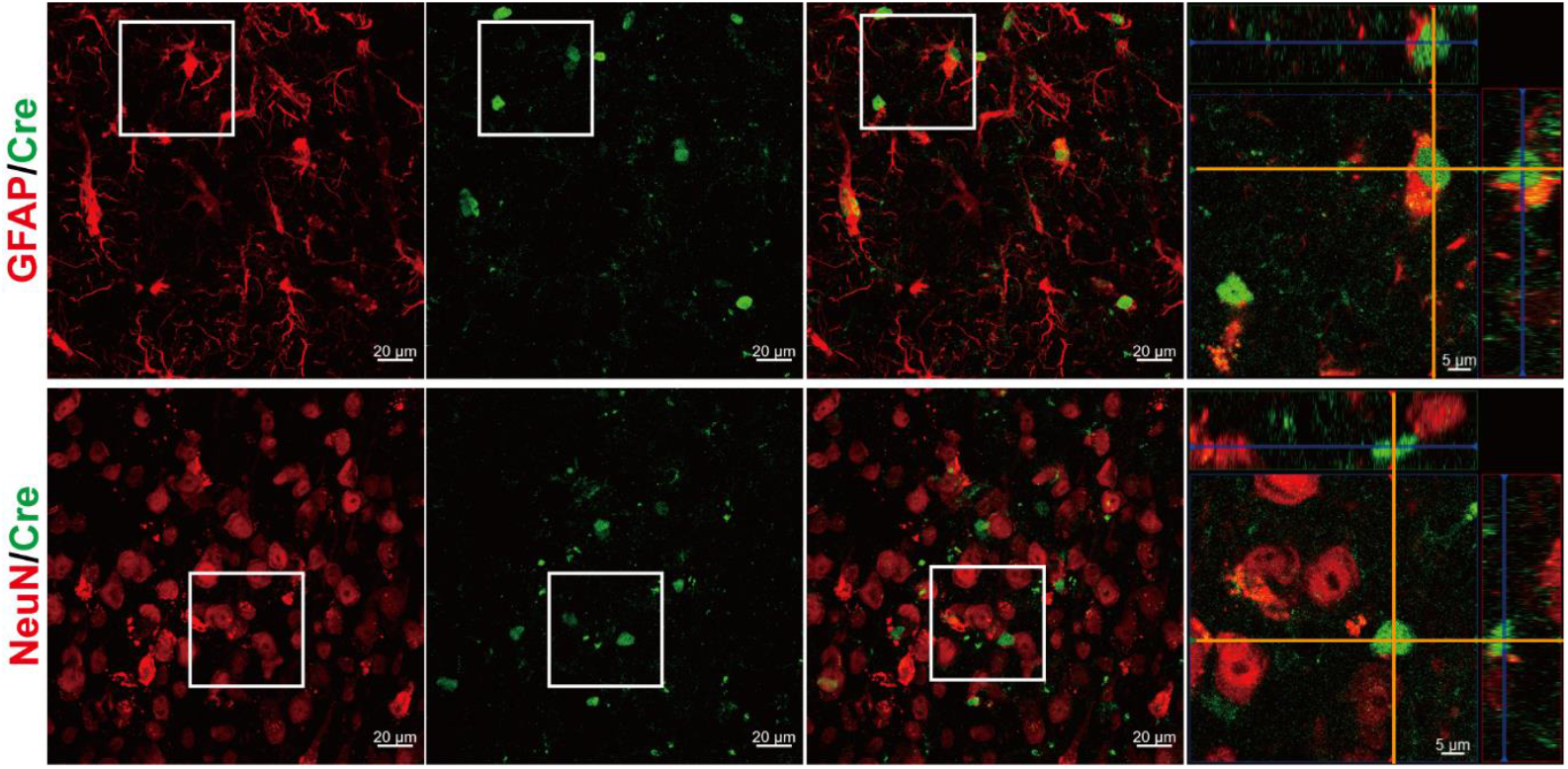
AAV9 GFAP::Cre expression in GFAP^+^-astrocytes but not in NeuN^+^-neurons in the mouse cortex. AAV dosing at 1×10^10^ gc/ml, 1 μl; 14 days post viral injection (dpi).

### Astrocyte-to-Neuron conversion in Aldh1l1-CreER^T2^ mice

Wang et al (51) reported that when Aldh1l1-CreER^T2^ mice were crossed with R26R-YFP mice to label some of the astrocytes with YFP after administration of tamoxifen, the YFP-labeled astrocytes were difficult to convert into neurons. Our group has performed similar lineage tracing experiments but observed clear astrocyte-to-neuron conversion (Fig. 2-3). We crossed Aldh1l1-CreER^T2^ mice with Ai14 mice and administered tamoxifen to induce Cre-mediated recombination so that some of the astrocytes will be labeled by tdTomato (Fig. 2). As expected, the tdTomato-labeled cells were immunopositive for astrocyte marker GFAP/S100β in non-viral infected cortex (contralateral to the viral injected hemisphere) (Fig. 2A left panel, and Fig. 2B). In contrast, in AAV5 GFAP::NeuroD1-infected cortex, some of the tdTomato-labeled cells lost GFAP/S100β signal, and displayed typical neuronal morphology (Fig. 2A right panel, and Fig. 2C). Further immunostaining with neuronal marker NeuN confirmed the neuronal identity of some of the tdTomato-labeled cells in NeuroD1-infected cortex (Fig. 3). Therefore, these astrocytic lineage tracing experiments in Aldh1l1-CreER^T2^ mice clearly demonstrate that astrocytes can be directly converted into neurons by NeuroD1, consistent with our series of publications in recent years (31–33, 59–62).

**Fig. 2.**
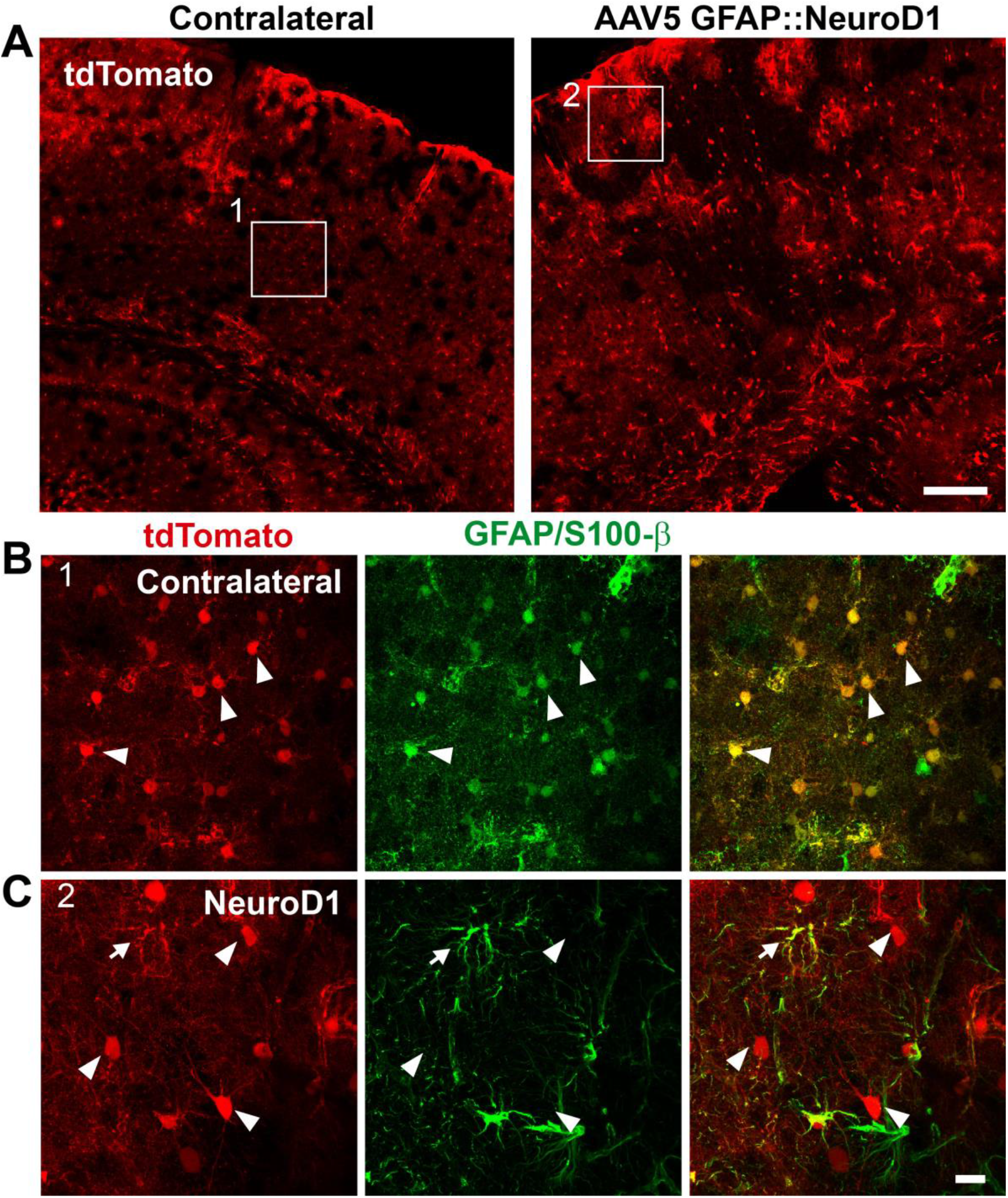
Lineage traced astrocytes in Aldh1l1-CreER^T2^ mice crossed with Ai14 mice. A, tdTomato-labeled cells in AAV5 NeuroD1-infected cortex (right panel) and non-infected contralateral cortex (left panel). B, In contralateral side without viral injection (Box1 in panel A), tdTomato-labeled cells were GFAP/S100β-positive astrocytes, as expected. C, In NeuroD1-infected cortex (Box2 in panel A), some tdTomato-labeled cells showed clear neuronal morphology and not co-localized with GFAP/S100β.

**Fig. 3.**
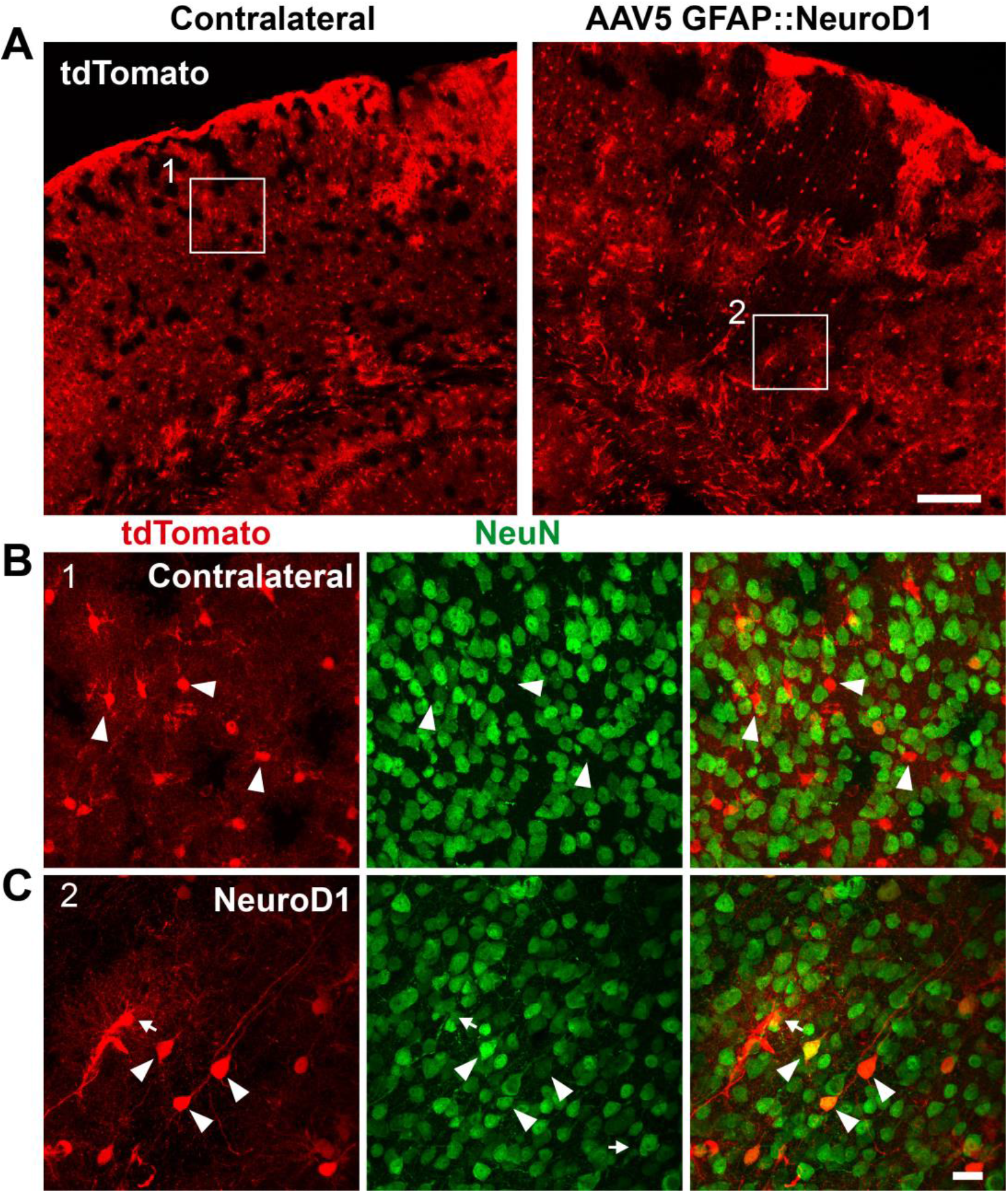
tdTomato-labeled astrocytes in Aldh1l1-CreER^T2^ mice converted into NeuN^+^ neurons after infected by AAV5 GFAP::NeuroD1. A, tdTomato-labeled cells in non-infected contralateral cortex (left panel) and NeuroD1-infected cortex (right panel). B, In non-infected contralateral side (Box1 in panel A), tdTomato-labeled cells were rarely colocalizing with NeuN. C, In NeuroD1-infected cortex (Box2 in panel A), some tdTomato-labeled cells were co-localized with NeuN, indicating that they have been converted into neurons. Note that the number of tdTomato-labeled astrocytes decreased significantly in the NeuroD1-converted areas, further suggesting that these tdTomato-labeled neurons were originally converted from tdTomato-traced astrocytes.

### Neuronal conversion induced by retrovirus overexpressing NeuroD1 in dividing glial cells

While AAV has the advantage of low immunogenicity and relatively safe as a gene therapy vector for the treatment of neurological disorders, its capability to infect both neurons and glial cells may cause confusion if AAV dosing and promoter are not handled properly. Therefore, if one’s main research purpose is not to generate as many neurons as possible to treat certain neurological disorders, retroviruses that mainly target dividing glial cells may be a better choice to study basic molecular mechanisms of glia-to-neuron conversion. We have previously reported that retroviruses expressing NeuroD1 can convert dividing glial cells into neurons (31, 32). Here, we provide another example of using retrovirus, instead of AAV, to ectopically express NeuroD1 in dividing glial cells and convert glial cells into neurons (Fig. 4). When injecting retroviruses into adult mouse cortex, because neurons cannot divide and therefore retroviruses cannot enter neuronal nuclei, only dividing glial cells can allow retroviruses enter glial nuclei to express transgene. Therefore, retroviruses should always be readily deployed if any confusion arises regarding AAV results. To conclude, if someone still has any doubt on whether certain transcription factor(s) can convert glial cells into neurons or not, then using retrovirus to express the transgene(s) should be a safe way to unambiguously test glia-to-neuron conversion without the complication of AAV.

**Fig. 4.**
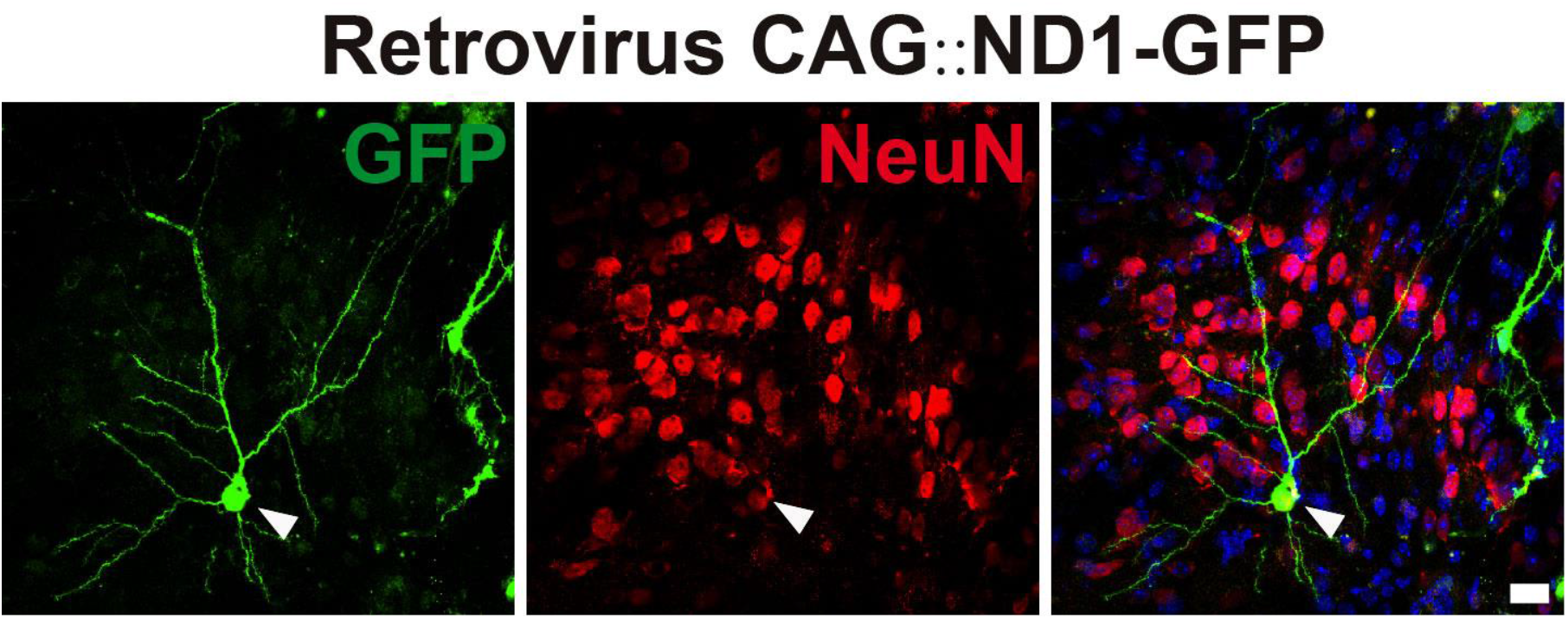
Retrovirus as an important tool to target dividing glial cells more specifically than AAV. Ectopic expression of NeuroD1 through retroviruses (CAG::NeuroD1-GFP, 1×10^7^ gc/ml, 1 μl) in the dividing glial cells of the mouse cortex converted glial cells into neurons (14 dpi). For more retrovirus info, see Guo et al (31).

## DISCUSSION

In recent years, many groups have used AAV-mediated ectopic expression of transcription factors or knockdown of PTBP1 to convert resident glial cells into neurons. However, Wang et al. (51) used a rather high dosage of AAV (10-1000 folds higher than that used in our lab or other labs) to challenge the field of *in vivo* reprogramming. In this responding article, we point out that the high dosage of AAV used by Wang et al (51) is destined to produce artifacts, as shown by their GFAP::Cre expression in neurons instead of astrocytes. We also provide further evidence to demonstrate unambiguously that glial cells can be converted into neurons by ectopic expression of NeuroD1 through lineage tracing or retroviral expression experiments.

Given such artifacts arising from a prominent lab, we feel that it is important to lay out some principles regarding how to make a right judgement on genuine *in vivo* glia-to-neuron conversion:

**First,** one must take a wholistic view on the entire *in vivo* glia-to-neuron conversion field before focusing on one single experiment, which can be an artifact produced by a specific person.
**Second,** one must test different doses of the delivery vehicles (viral or non-viral) to find optimal dosing for certain experiments. In particular, the toxic effects of high dosing should be tested because it is obvious that our brain cannot tolerate a huge amount of viral infection (or non-viral particles).
**Third,** for any factor(s) that is claimed to be capable of converting glial cells into neurons, we recommend conducting both *in vitro* and *in vivo* studies and use both retrovirus and AAV (or lentivirus) to unambiguously demonstrate the glia-to-neuron conversion. <u>Note that, AAV is great for *in vivo* work but infects cultured astrocytes with relatively low efficiency. Retroviruses are better for *in vitro* cultured astrocytes</u>.
**Last but not least,** for anyone who has benefit of doubt on *in vivo* glia-to-neuron conversion, please keep your mind open. Make comments specific on the data, and don’t simply criticize new discoveries using “I can’t believe” as a non-scientific argument. After all, scientific new discoveries are pushing the boundaries of our understanding every day.

### Virus dosing is critical to avoid artifacts

Injecting high dosage of AAV into the brain may result in some artifacts that are very difficult to interpret. Wang et al (51) injected 1 μl of 2×10^13^ GC/ml AAV5 GFAP::Cre into the mouse cortex and found that the Cre signal was predominantly detected in neurons instead of astrocytes. It is certainly difficult to understand why their Cre expression was found in neurons under the control of astrocytic promoter GFAP. In their discussion part, they attributed this phenomenon to exosomes or tunneling nanotubes induced by some uncertain genetic manipulations (51). While this is one possibility, a more straightforward explanation is the toxic effects to neurons caused by high dosage of virus administration. In our previous studies, we proved that 1-2×10^10-11^ of AAV GFAP::Cre (only 1/100 to 1/1000 of their dosage) was sufficient to trigger Cre-mediated recombination in the mouse cortex and striatum (32, 33). Most importantly, as repeated here in this study, Cre expression was restricted to astrocytes at this low dose. It is incomprehensive why Wang et al (51) applied 1000-fold higher dosage of GFAP::Cre without questioning their own data of Cre expression in neurons. They should have investigated immediately why Cre was mostly expressed in neurons, and by lowering AAV dosage they could have found the right answer quickly without falsefully challenging the field of *in vivo* reprogramming based on one set of improperly designed experiments.

### NeuroD1-induced astrocyte-to-neuron conversion through lineage tracing

Wang et al (51) reported their lineage tracing experiments by crossing tamoxifen-inducible *Aldh1l1-CreER*^*T2*^ transgenic mice with a reporter line (*R26R-YFP or Ai14*) to trace astrocytes labeled by YFP. We have conducted almost the same experiments using *Aldh1l1-CreER*^*T2*^ mice crossed with a different reporter line Ai14. Surprisingly, while we report here clear astrocyte-to-neuron conversion through astrocyte lineage-tracing experiments, Wang et al (51) reached opposite conclusion of not detecting NeuN^+^ neurons. Comparison of the two studies identified immediately the time difference of the results reported after NeuroD1 AAV injection: we found clear conversion of tdTomato-traced astrocytes into neurons at 135 days post AAV NeuroD1 injection (experiment delayed by COVID-19); while Wang et al (51) stopped short of their experiments at 28 days post AAV NeuroD1 injection. We have already informed the senior author of Wang et al (51) (C-L Zhang) about our lineage tracing results, and they promised to observe longer time in their lineage tracing experiments. We look forward to hearing from them soon and seeing their updated version of the article, hopefully together with their lowered dosage of GFAP::Cre results. In fact, even in the present data of Wang et al (51), the morphology of the NeuroD1-infected YFP-traced astrocytes was obviously different from that of the control group. In their NeuroD1 group (see Wang et al, Fig. 5F and Fig. 6F), the NeuroD1-infected YFP-traced astrocytes displayed clear morphological changes toward neuronal like structures with many fine processes already retracted in comparison to the control group. It is rather astonishing that the authors of Wang et al (51) would ignore such evident morphological changes and abruptly ended their experiments at 28 days post NeuroD1 AAV infection. We sincerely hope that Wang et all will soon provide longer time point data to tell the world whether those morphologically changed YFP-traced astrocytes will become NeuN^+^ neurons or not.

### How to interpret the BrdU data in a right way?

Quiescent or resting astrocytes are more resistant to cell conversion compared to reactive astrocytes, which explains why previous studies targeted more on reactive astrocytes for *in vivo* reprogramming (31–33, 36, 43, 44, 49, 50, 63–65). However, Wang et al (51) used BrdU-incorporation experiment to declare that converted neurons were not derived from BrdU^+^ reactive astrocytes, largely due to their poorly designed experiments for BrdU-labeling. The major flaw of their BrdU experiment is that they have administered BrdU for such a long-time span of weeks after AAV NeuroD1 injection, leading to a large number of BrdU^+^ astrocytes that have never had a chance to be infected by AAV NeuroD1. Therefore, they of course could not detect many BrdU^+^ neurons and the ratio of BrdU^+^NeuN^+^ neurons was artificially low among all the BrdU^+^ cells. The right experiment should be to inject AAV NeuroD1 at the end of their BrdU labeling in order to convert many BrdU-labeled astrocytes into neurons. BrdU labeling should be stopped immediately after AAV injection to prevent further BrdU-labeling.

### How to understand the puzzle of neuronal density not changed after conversion?

Wang et al (51) was puzzled by the fact that after astrocyte-to-neuron conversion, there was no significant increase of neuronal density. We have essentially observed the same phenomenon in non-injured non-diseased mouse brains. However, in injured brains with substantial neuronal loss, we always detect a significant increase of neuronal density across the entire injury/diseased areas. In fact, from the data presented by Wang et al (51), the tissue repair is so obvious in their Fig. 2 (C, E) and their Fig. 6F, as shown by significantly reduced cortical tissue loss in the NeuroD1 group compared to their control group, which is also consistent with our reported findings (61). Wang et al (51) ignored the apparent tissue repair in the center of lesion core in the NeuroD1 group, and asked why neuronal density did not increase significantly in the less injured surrounding areas. This is actually similar to our findings in the mouse striatum of Huntington’s disease model where the neuronal density did not change much after conversion but the overall striatal atrophy was alleviated (33). Wang et al (51) assumed that the neuronal density should increase after conversion, but they probably did not realize that their highly toxic AAV dosage already damaged many neurons, eventually leading to a balance between the newly converted neurons and the loss of preexisting neurons. We hypothesize that there should be some kind of homeostatic control to keep the neuron density in certain brain regions relatively constant to maintain normal functions, which surely warrants further studies.

Besides neuronal density, there is also some concern in the field that astrocyte-to-neuron conversion might lead to the depletion of astrocytes in the converted areas. Fortunately, we have never observed any depletion of astrocytes in NeuroD1-converted areas in mouse, rat, and monkey brains. In fact, the results from Wang et al (51) confirmed our observations that astrocytes were not depleted in NeuroD1-expressed areas, consistent with the notion that astrocytes are dividing cells with proliferative capability (66, 67). Our recent study detected more proliferative astrocytes (Ki67+) in the converted areas, indicating that astrocytes can repopulate themselves after some of the astrocytes being converted into neurons (33)(61).

### Recommendation for future research

Given the fact that C-L Zhang’s lab was among the early pioneers who reported *in vivo* glia-to-neuron conversion, the impact of the pre-print article of Wang et al (51) would pose grave danger to the field of *in vivo* reprogramming if their flawed design and wrong interpretations were not corrected immediately. While it is up to every single scientist to make his or her own judgement, <u>we do want to reiterate the importance of using different dose, different types of viral vectors, and perform both *in vitro* and *in vivo* experiments to prove or disprove any hypothesis</u>. We do have every reason to believe that Wang et al (51) might have good intention to raise a potential problem to the field, but such hasty deposit of improperly designed experiments based solely on one single high dosing of AAV without verification by retrovirus and *in vitro* studies, should be highly discouraged in future studies.

## Materials and Methods

### Mouse

8-10-week-old mice were used in this experiment. The wildtype C56BL/6J mice were purchased from Guangdong Medical Laboratory Animal Center (Guangzhou, China), Aldh1l1-Cre^ERT2^ transgenic mice (031008) and Ai14 knock in mice (#007914) were from Jackson Laboratory. All animals were housed in a 12 h light/dark cycle and supplied with sufficient food and water. All the experiments were approved by Jinan University laboratory animal ethics committee.

### Virus information

Single strand adenovirus-associated viral (ssAAV, AAV for short) vector hGFAP::Cre and FLEX-CAG::mCherry were constructed as previously described (32), and used for Cre experiment (Fig. 1). A short version of hGFAP promoter (681 bp) was also used in this study (68) for the lineage tracing experiment (Fig. 2-3). AAV serotype 9 (AAV9) and 5 (AAV5) were produced by PackGene^®^ Biotech, LLC, purified through iodixanol gradient ultracentrifuge and subsequent concentration. Purified AAV viruses were tittered using a quantitative PCR-based method. All AAV used in this study was prepared in 0.001% Pluronic F-68 solution (Poloxamer 188 Solution, PFL01-100ML, Caisson Laboratories, Smithfield, UT, USA). Retroviral vector CAG::NeuroD1-IRES-GFP were constructed, packaged and concentrated as previously described (32) for the retrovirus experiment (Fig. 4).

### Mouse Model of Ischemic Injury and Virus Injection

Endothelin-1 (ET-1, 1-31) was injected into motor cortex of the adult WT C56BL/6J mice to create a focal ischemic injury as described (32), for the Cre experiment. Briefly, the mice were anesthetized with 20 mg/kg 1.25% Avertin (a mixture of 12.5 mg/mL of 2,2,2-Tribromoethanol and 25 μL/mL 2-Methyl-2-butanol, Sigma, St. Louis, MO, USA) through intraperitoneal injection and then placed in a prone position in the stereotaxic frame. 1 μL of ET-1 (1 μg/μL dissolved in PBS) was injected at the following coordinate: +0.2 mm anterior-posterior (AP), ± 1.5 mm medial-lateral (ML), 1.2 mm dorsal-lateral (DV) at the speed of 100 nl/min. After injection, the pipette was kept in place for about 10 minutes and then slowly withdrawn. 7 days later, 1 μL of virus mixture AAV9 hGFAP::Cre (1×10^10^ GC/ml) and FLEX-CAG::mCherry (1×10^12^ GC/ml) was injected at the same coordinates.

For intact mouse cortex, 1 μL of retroviruses CAG::NeuroD1-IRES-GFP (1×10^7^ TU/ml) or 1 μL of AAV5 GFAP::NeuroD1 (1×10^12^ GC/ml) were injected at the similar coordinates described above.

### Immunofluorescence

The mice were anesthetized with 2.5% Avertin and then sequentially perfused intracardially first with saline solution (0.9% NaCl) and then with 4% paraformaldehyde (PFA). The brains were collected and post-fixed in 4% PFA overnight and sequentially placed in 20% and 30% sucrose at 4°C until the tissue sank. The dehydrated brains were embeded in Optimal Cutting Temperature (Tissue-Tek® O.C.T. Compound, Sakura® Finetek, Torrance, CA, USA), and then serially sectioned at the coronal plane on the cryostat (Thermo Scientific, Shanghai, China) at 30 μm thickness. For immunofluorescence, free floating brain sections were first washed with PBS and blocked for 1 hour at room temperature (RT) in 5% normal donkey serum, 3% bovine serum albumin and 0.3% TritonX-100 prepared in PBS, and then incubated overnight at 4 °C with primary antibodies diluted in blocking solution. After additional washing with 0.2% PBST (0.2% tween-20 in PBS), the samples were incubated with 4′,6-diamidino-2-phenylindole (DAPI; F. Hoffmann-La Roche, Natley, NJ, USA) and appropriate donkey anti-mouse/rabbit/rat/chicken secondary antibodies conjugated to Alexa Fluor 488, Alexa Fluor 555, or Alexa Fluor 647 (1:1000, Life technologies, Carlsbad, CA, USA) for 2 hours at RT, followed by extensive washing with PBS. Samples were finally mounted with VECTASHIELD® mounting medium (VECTOR Laboratories, Burlingame, CA, USA) and sealed with nail polish. Representative Images were taken with confocal microscope (LSM880, Zeiss, Jena, Germany).

Primary antibodies used were listed as follows: rabbit anti-GFAP (a marker for astrocytes, 1:1000, Cat# Z0334, DAKO), rabbit anti-NeuN (a marker for neurons 1:1000, Cat# ab177487, Abcam, Cambridge, Massachusetts, USA), rabbit anti-S100β (a marker for astrocytes, 1:500, Cat# ab52642, Abcam), mouse anti-Cre recombinase (1:500, Cat# MAB3120, Millipore), chicken anti mCherry (1:1000, Cat# ab205402, Abcam).

